# Electron microscopy mapping of the DNA-binding sites of monomeric, dimeric, and multimeric KSHV RTA protein

**DOI:** 10.1101/2023.05.01.538939

**Authors:** Jayla C. Calhoun, Blossom Damania, Jack D. Griffith, Lindsey M. Costantini

**Affiliations:** Biological and Biomedical Sciences Department, North Carolina Central University, Durham, North Carolina, USA; Lineberger Comprehensive Cancer Center, University of North Carolina, Chapel Hill, North Carolina, USA

## Abstract

Molecular interactions between viral DNA and viral-encoded protein are a prerequisite for successful herpesvirus replication and production of new infectious virions. Here, we examined how the essential Kaposi’s sarcoma-associated herpesvirus (KSHV) protein, RTA, binds to viral DNA using transmission electron microscopy (TEM). Previous studies using gel-based approaches to characterize RTA binding are important for studying the predominant form(s) of RTA within a population and identifying the DNA sequences that RTA binds with high affinity. However, using TEM we were able to examine individual protein-DNA complexes and capture the various oligomeric states of RTA when bound to DNA. Hundreds of images of individual DNA and protein molecules were collected and then quantified to map the DNA binding positions of RTA bound to the two KSHV lytic origins of replication encoded within the KSHV genome. The relative size of RTA or RTA bound to DNA were then compared to protein standards to determine whether RTA complexed with DNA was monomeric, dimeric, or formed larger oligomeric structures. We successfully analyzed a highly heterogenous dataset and identified new binding sites for RTA. This provides direct evidence that RTA forms dimers and high order multimers when bound to KSHV origin of replication DNA sequences. This work expands our understanding of RTA binding, and demonstrates the importance of employing methodologies that can characterize highly heterogenic populations of proteins.

**Importance:** Kaposi’s sarcoma-associated herpesvirus (KSHV) is a human herpesvirus associated with several human cancers, typically in patients with compromised immune systems. Herpesviruses establish lifelong infections in hosts in part due to the two phases of infection: the dormant and active phases. Effective antiviral treatments to prevent the production of new viruses are needed to treat KSHV. A detailed microscopy-based investigation of the molecular interactions between viral protein and viral DNA revealed how protein-protein interactions play a role in DNA binding specificity. This analysis will lead to a more in depth understanding of KSHV DNA replication and serve as the basis for anti-viral therapies that disrupt and prevent the protein-DNA interactions, thereby decreasing spread to new hosts.

## Introduction

Our model for studying viral DNA-binding proteins is one of the eight known human herpesviruses, Kaposi’s Sarcoma Associated Herpesvirus (KSHV, HHV-8). KSHV is the etiological agent responsible for several human diseases including Kaposi’s sarcoma (KS), Multicentric Castleman’s Disease (MCD), Primary Effusion Lymphomas (PEL) and KSHV inflammatory cytokine syndrome (1–4). Drugs currently used to treat KSHV associated malignancies are at best moderately effective and therapies for related human herpesviruses have been shown to be ineffective (5–9). A specific cure or vaccine for the treatment or prevention of KSHV has not yet been approved for clinical use (5,6,10). Given the endemic spread of the disease in Africa, and prevalence in transplant and HIV-infected patients, there is still a need for novel KSHV therapeutic targets (5,9). Herpesviruses contain a double stranded DNA genome and encode their own DNA replication proteins that are essential for successful viral production and dissemination. Therefore, it is critical to characterize the protein-DNA interactions that contribute to viral proliferation, with the goal being to potentially disrupt them and prevent viral spread.

Like all members of the *Herpesviridae* family, KSHV has a biphasic life cycle, consisting of a prolonged latency with reoccurring episodes of lytic reactivation. Both phases contribute to the pathogenesis of disease and promote lifelong infections of hosts (11,12). However, it is only during the KSHV lytic cycle that new infectious virions are produced. A key step to generating new virions is successful viral DNA replication. Latent viral DNA replication relies on host-cell proteins, while lytic viral DNA replication is controlled by the KSHV encoded viral DNA replication proteins (13). In this study, we focused on one of the essential KSHV DNA replication proteins, RTA (replication and transcription activator) encoded by *ORF50* (14). RTA is known to have dual roles in lytic reactivation, including (i) activating downstream lytic genes as a viral transcription factor (15,16) and (ii) binding the KSHV origin of replication DNA as an origin binding protein (17–19). Moreover, it has previously been demonstrated that RTA expression is sufficient and necessary to induce reactivation of the lytic phase in various KSHV cell culture model systems (20–22). Thus, RTA is an appealing anti-viral therapeutic target, due to its key activity as a lytic switch protein (23,24) and role in initiating of lytic DNA replication.

The cumulative results of previous studies characterizing RTA have shown that RTA expression is highly regulated and RTA forms complexes with host (24–27) and viral (27,28) proteins (19,29–31). Traditional, gel-based studies with truncated forms of RTA (C-terminus deletion, 1-321 amino acids, aa), full-length RTA (1-691 aa) and RTA with internal deletions, identified the DNA binding domain (1-390 aa), the DNA binding inhibitory sequence (490-535 aa) and the dimerization domain (1-414 aa) (31,32). Accordingly, studies have also been used to identify interaction domains and characterize cooperativity between RTA and other trans-acting cellular factors (MDM2, OCT1, RBJ-k, etc.) that have been shown to regulate abundance of the protein, mediate RTA autoregulation and activate cellular pathways during reactivation (28,31– 34). The expected monomeric molecular weight of RTA is ∼74 kDa, however, the addition of posttranslational modifications increases the predicted molecular weight to approximately ∼110 kDa (35). It has also been suggested that the protein does not function as a monomer, but forms tetramers and higher order multimers when binding DNA (32). Experiments combining full-length RTA and a C-terminus deletion mutant showed inhibition of RTA binding to viral promoter regions, further demonstrating that the ability of RTA to form homodimers/multimers may be crucial for its activity (20). Overall, our understanding of the observed molecular weight and oligomeric state of RTA has been informed by traditional assays (immunoblotting, affinity-IP, EMSA) that look at the predominant protein species (32,35,36).

The previously identified RTA-DNA binding sites are known as RTA response elements or RREs (37). RTA, as a potent transactivator, binds to RRE in both early and late lytic viral promoters (38). As a DNA replication protein, RTA binds to sequences within the two KSHV lytic origins of replication (OriLyt-L and -R), which share a ∼1.2 kb long homologous sequence (39). RTA binding to the KSHV lytic origins initiates DNA replication by recruiting the core DNA viral replication proteins: *ORF9*/polymerase, *ORF59*/polymerase processivity factor, *ORF6*/single-stranded binding protein, *ORF56*/primase, *ORF44*/helicase, and *ORF40/41*/primase associated factor (40). And it has been determined that the 32 bp RTA response element (5’ CTACCCCCAACTGTATTCAACCCTCCTTTGTTT 3′) found in origin DNA is required for OriLyt dependent DNA replication (19,39,41). These previous studies form the foundation of our understanding about RTA.

In this study, we directly visualized purified viral proteins and viral DNA via TEM to characterize the molecular interactions involved in initiating DNA replication for a human herpesvirus. We also expanded upon our understanding of the degree to which DNA binding locations and protein conformation are heterogenous and how protein monomers and dimers function differently with respect to DNA-binding location specificity. TEM images of individual DNA and protein molecules were collected and then quantified to map the DNA binding positions of RTA bound to the KSHV OriLyt-L and -R and the relative size of RTA unbound or bound to DNA were compared to protein standards (42,43) to ascertain whether RTA was monomeric, dimeric, or oligomeric under the different conditions tested. Using our TEM approach to directly visualize RTA and KSHV DNA, we successfully quantified a highly heterogenous dataset and identified new binding sites for RTA and provide evidence that suggests RTA forms dimers and high order multimers when bound to OriLyt DNA. This study enhances our understanding of human herpesvirus DNA replication proteins and more specifically, we show that the viral origin binding protein binds to discrete locations within the viral origin DNA and the binding sites are distinctive for the protein monomers and dimers.

## Materials and Methods

OriLyt-L was isolated from pDA15 (44) a generous gift from David AuCoin and Cyprian Rossetta at the University of Nevada. OriLyt-R DNA was synthesized using BAC16 KSHV sequence (accession MK733609) via GeneArt (ThermoFisher) and subcloned into pRSET-A plasmid via NdeI/XhoI, see Supplemental Table S1 primer pair 1. To purify the 1.8 kb OriLyt-L fragment of DNA, the plasmid containing the origin DNA sequence was digested with HindIII and EcoRI (New England Biolabs) according to manufacturer’s protocol (Supplemental Figure S2A, B). To isolate the approximately 2.4 kilobase (kb) OriLyt-R fragment, pRSET-A-OriLyt-R was digested with XhoI, ScaI and NdeI (New England Biolabs) according to manufacturer’s protocol (Supplemental Figure S2C, D). DNA fragments were separated using 0.8% agarose gel and then purified using Qiaquick gel extraction kit (Qiagen). To incorporate biotin at the 5’ end of OriLyt DNA, digested fragments were incubated with Klenow Exo-polymerase (New England Biolabs) with 2.8µM biotin-dCTP (Invitrogen) for 1 hour at 37° C prior to gel extraction.

For lytic origin DNA fragments A-C, PCR amplification of the pDA25 OriLyt-L or pRSET-A-OriLyt-R using primer pairs 2-6 (Supplemental Table S1). Either AccuPrime Pfx DNA polymerase (Invitrogen) or Platinum Taq DNA polymerase (Invitrogen) were used, per the manufacturer’s instructions, to amplify viral DNA fragments. Fragments were then isolated by gel extraction. For AP-1 site fragments, single-stranded DNA oligos (Supplemental Table S2) were synthesized (Eurofins) and annealed.

Full length (aa 1-691) and truncated (aa 1-321) RTA sequences were expressed and purified using SF9 insect cell expression system (ThermoFisher). Gene sequences were based on JSC-1 sequence (accession GQ994935). Alcohol dehydrogenase (Sigma) and conalbumin (Cytiva, Gel Filtration Calibration Kits) were commercially purchased and resuspended using manufacture’s recommendations.

### Preparation of DNA and protein for transmission electron microscopy (TEM)

DNA-protein binding assays were conducted at a mass ratio of 2:1 (DNA: RTA) in 50 or 100 ul total reaction volume (4mM HEPES, 10mM NaCl, 0.1mM DTT, 0.1mM EDTA) for 30 or 60 minutes at room temperature. Briefly, 400 ng of linearized DNA (OriLyt-L, -R) was incubated in the reaction buffer at room temperature for 10 mins. Next, RTA (200 or 400 ng) was added to the reaction with DNA and incubated for an additional 30 minutes at room temperature. To label the 5’ biotin DNA with streptavidin, 0.01mg/ml streptavidin (Invitrogen) was added to binding reaction and incubated for 20 mins at room temp. Samples were either added to size exclusion column directly or fixed with 0.6% glutaraldehyde for 5 mins at room temperature and passed over a 2 ml column of 2% agarose beads (Agarose Bead Technologies) equilibrated in 10mM Tris-HCl (pH 7.6) and 1.0 or 0.1 mM EDTA. Sample fractions were collected for direct mounting onto carbon supports for tungsten shadow casting and TEM(45). For individual proteins, RTA, truncated RTA, conalbumin and alcohol dehydrogenase were diluted in 4mM HEPES, 10mM NaCl, 0.1mM DTT, 0.1mM EDTA and prepared for TEM.

### Tungsten-shadow casting

DNA or DNA-protein complexes were prepared by tungsten rotary shadow casting as previously described (46). Sample fractions were mixed with 2mM spermidine and incubated on glow discharge treated charged carbon coated 400-copper mesh grids (Electron microscopy sciences, EMS) for 3 minutes. Carbon grids were washed in high grade distilled water and dehydrated in a series of ethanol washes (25, 50, 75, 100%), air-dried, and rotary shadow-cast with tungsten (EMS). Samples were visualized on a FEI Technai 12 or Philips CM12 TEM at 40kV. Transmission electron microscopy (TEM) images were captured on a Gatan First Light CCD camera using Gatan Digital Micrograph software (Gatan, Pleasanton, CA). TEM micrographs were contrast adjusted and inverted using Adobe Photoshop software.

### Electrophoretic Mobility Shift Assay (EMSA)

For EMSA binding reactions, 12ug of RTA and 50-80ng DNA were incubated in 50 mM NaCl, 20 mM HEPES, 0.1 mM EDTA, 0.5 mM DTT, 13% glycerol in a total reaction volume of 20 ul. Reactions were incubated on ice for 30 minutes. For competition assays, unlabeled competitor DNA (200-300 ng) was added in 4-5-fold excess to the reaction prior to the specific DNA. Reactions were loaded into a 1x TAE, 0.7-1.0% agarose gel (Invitrogen), pH 9.15, and run for 90 minutes at 75 volts then 60-90 minutes at 90 volts. Gels were imaged using BioRad Chemidoc system (BioRad) and DNA was visualized as total DNA via gel red staining (Biotium) or using oligo specific dye (Alexa-488, Cy-5 or Cy-3).

### Data acquisition and analysis

Distances along the length of the DNA, and area of protein were manually traced using Gatan Digital Micrograph or ImageJ software (NIH). Data sets for length and area were measured in pixels and pixels^2^, respectively. To convert pixels to nm, data was multiplied by a conversion factor derived from the 200 nm scale bar visible in all collected TEM images divided by its length measured in pixels. To convert nm to base pairs, DNA measurements in nm were multiplied by the known length of base pairs in nm (47,48).

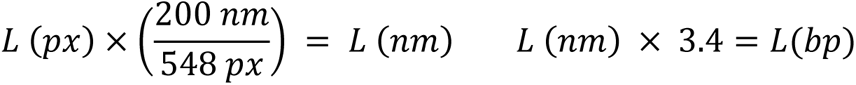

Following quantification of all the parameters for each DNA-protein complex, the data was analyzed using Microsoft excel, Prism software (GraphPad), and sequence specific analysis was completed using CLC Main WorkBench20 (Qiagen).

## Results

### TEM data reveals heterogeneity of DNA-protein binding of KSHV RTA to lytic origin DNA

We visualized KSHV origin binding protein, RTA, in the presence of circular or linear OriLyt DNA at nanometer resolution by using specialized transmission electron microscopy (TEM). Purified DNA and protein were incubated in solution and absorbed on to the surface of carbon coated copper mesh TEM grids and rotary shadow-cast with tungsten (46). This method permits imaging of proteins and DNA molecules and was successfully used to capture both unbound RTA (arrows) and RTA bound (arrowhead) to plasmid DNA containing the KSHV lytic origin (OriLyt) DNA sequence and isolated OriLyt linear DNA (Fig 1A, Supplemental Fig S1A-B). The KSHV genome adopts different conformations; a circular episome during latency and linear within infectious virions, while the DNA architecture during lytic genome replication is still widely unknown (49,50). Our method provides the flexibility of examining multiple DNAs at varying lengths. RTA associated with OriLyt DNA in both plasmid (supercoiled and relaxed) and linear conformations (Fig 1A, Supplemental Fig S1B, D). In parallel, we completed traditional electrophoresis mobility shift assays (EMSA) with RTA and the isolated left and right lytic origin DNA (OriLyt-L, ∼1.8kb and -R, ∼2.4kb). OriLyt-L and -R incubated in the presence of RTA shifted to a higher molecular weight band confirming the molecular interaction between protein and DNA via gel-based methods (Fig. 1B, lanes 1 and 7). These findings are consistent with previous studies (15), and provided the point of comparison for our TEM approach. To optimize the TEM sample preparation, add orientation to DNA, and enhance the population of RTA-DNA complexes while reducing the background of unbound proteins, additional steps were integrated into the methods pipeline (Supplemental Fig. S1C). These improvements helped reveal the heterogeneity of protein-DNA complexes, including RTA binding locations, protein dimensions, and DNA architecture (Fig. 1C and Supplemental Fig. S1D). Occasionally, we observed DNA formed loops at the location of RTA binding and more than one protein bound to an individual DNA (Supplemental Fig. S1D). Changes to DNA architecture and lower abundance molecular structures cannot be easily discerned using standard gel-based methods (Fig. 1B). Our observations from TEM indicated that RTA binds to multiple locations within the KSHV lytic origin sequences and binds to DNA in a variety of conformations.

**Figure 1.**
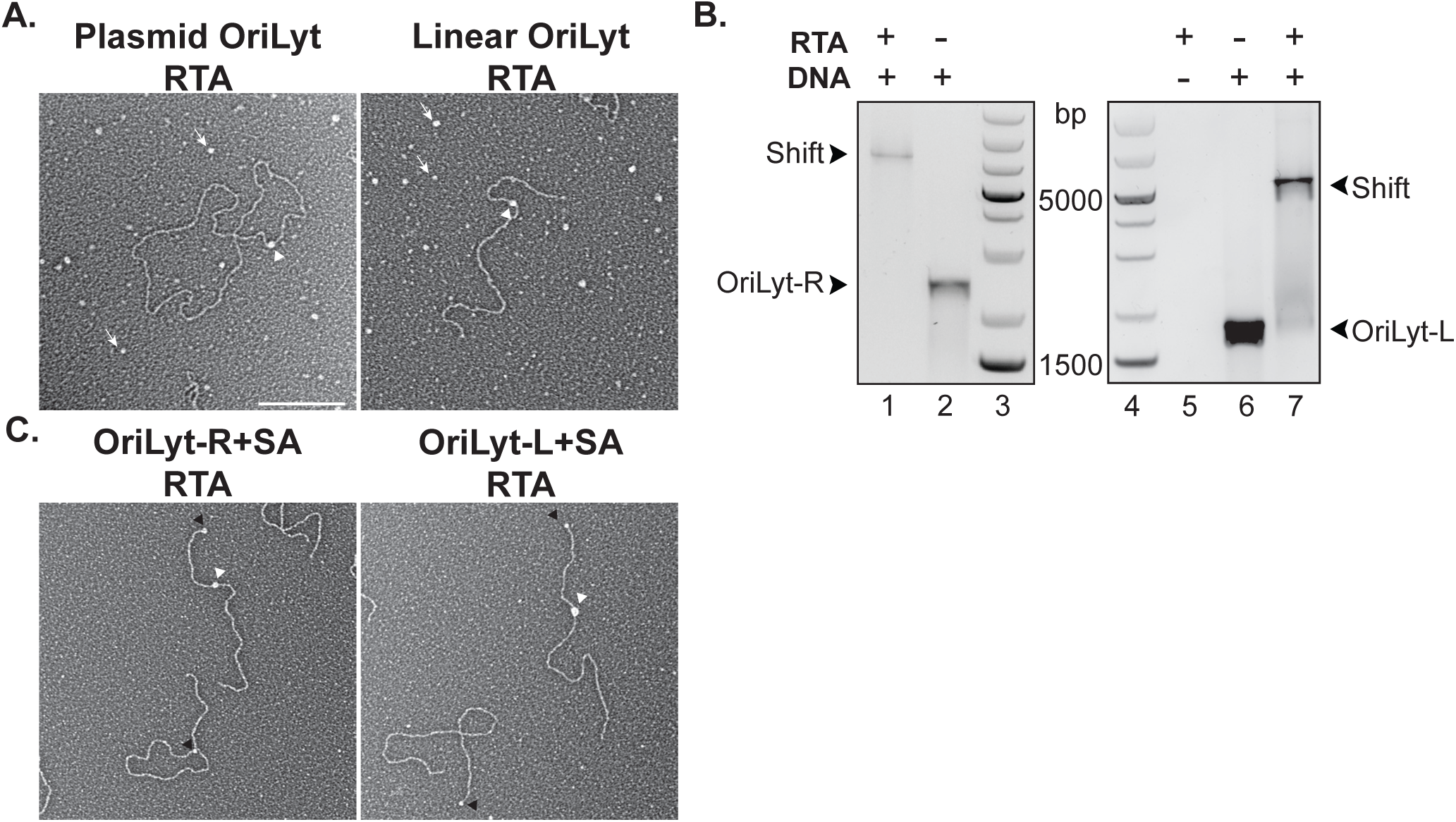
TEM and EMSA analysis of purified KSHV RTA and lytic DNA. A. Representative TEM micrographs of RTA (DNA and RTA, protein bound, white arrowhead; unbound protein, arrow) in the presence of plasmid containing OriLyt or isolated linearized OriLyt. B. Representative EMSA agarose gel of OriLyt-R and -L in the presence (Lanes 1, 7) or absence (Lanes 2, 6) of RTA. C. Representative TEM micrograph of RTA bound (white arrowhead) to streptavidin (black arrowhead), SA-end labeled OriLyt-L and OriLyt-R DNA. Scale bar=200nm.

### TEM mapping of RTA DNA-binding locations revealed specific DNA location preferences of RTA

To obtain protein positional data from TEM micrographs of RTA bound to streptavidin (SA) end-labeled DNA (Fig. 1C, 2A, B), we first preferentially labeled one end of the linearized OriLyt-L and -R (Supplemental Fig. 2) with biotin and SA and then measured the position of each DNA-bound RTA using three measurements: M1 full length of DNA, M2 distance from SA to RTA, and M3 distance from unlabeled DNA end to RTA (Fig. 2C). Table 1 summarizes the length measurements for all DNA molecules quantified (n=316 of OriLyt-L, n=270 of OriLyt-R). The average full length (M1) measured DNA length for OriLyt-L and -R were [556.2 +/- 30.45 nm] and [724.3 +/- 35.69 nm], respectively. The average protein footprint was determined by calculating the diameter of the protein area, assuming an average circular shape [16.4 +/- 3.12 nm] was used to set the histogram x-axis bin size (Fig. 2D,E). The y-axis corresponds to the number of RTA molecules located at a particular DNA site. For OriLyt-L, RTA has the highest tendency to bind 350 nm from the SA-labeled end of the DNA (Fig. 2B, D), while RTA had a greater propensity to OriLyt-R between 150-210 nm from the SA-labeled end of the DNA (Fig. 2C, E).

**Figure 2.**
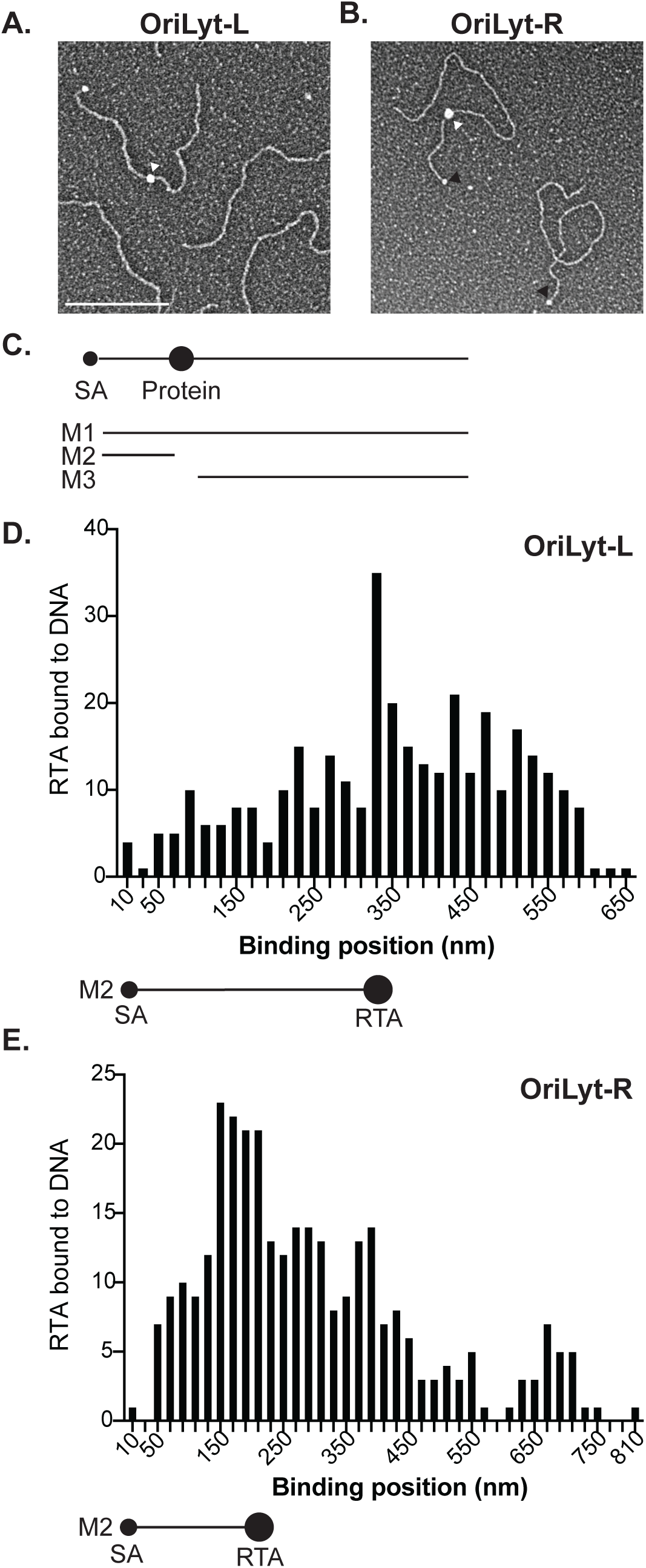
Mapping absolute binding position of RTA bound to KSHV lytic origin DNA. Representative TEM micrograph of RTA bound (white arrowhead) to SA-end labeled (black arrowhead) A. OriLyt-L or B. OriLyt-R C. Schematic of 3 length measurements (nanometers, nm), M1=full length of DNA, M2=streptavidin, SA to RTA, M3=Protein to unlabeled end of DNA. Histograms of the number of RTA molecules bound to KSHV OriLyt at the M2 distance (nm) D. Orilyt-L, n= 334 proteins or E. OriLyt-R n= 292 proteins. M2 schematic paired with histogram in D corresponds to RTA position (white arrowhead) in A. M2 schematic paired with histogram in E corresponds to RTA position (white arrowhead) in B. Scale bar=200nm.

**Table 1.** DNA length measurements for KSHV lytic origin DNA.

| **Table 1. DNA length measurements for KSHV lytic origin DNA** |  |  |  |
| --- | --- | --- | --- |
|  | Base pairs<br>*predicted* | M1 Mean +/-<br>SD (nm) | M1 (bp) |
| OriLyt-L (n= 316) | 1876 | 556.2 +/- 30.45 | 1871.2 +/- 105.8 |
| OriLyt-R (n= 270) | 2414 | 724.3 +/- 35.69 | 2436.5 +/- 119.9 |

Nanometer measurements were converted to base pairs (bp), to compare the corresponding binding location with the known DNA sequences (Table 1). The measured averages corresponded to the predicted lengths and gel analysis (Fig. 1). The average measured full length bp DNA length for OriLyt-L and -R were [1871.2 +/- 102.3] and [2436.5 +/- 119.9], respectively. Diagrams of OriLyt-L and -R (Fig. 3A, B) specify the relative base pair position of known TATA box (magenta), AP-1 (green), RTA RRE (purple) and A-T rich (cyan) regions (39). The most abundant measured binding site for RTA-OriLyt-L was between 900-1000 bp from the SA-labeled end of the DNA. This region coincides with the locations of an AP-1 site and a TATA box (Fig 3C, green and magenta lines). During sample preparation, a subset of experimental replicates were fixed with glutaraldehyde to preserve DNA-protein interactions. The total number of molecules (Fig. 3C, D) analyzed were divided into the two subgroups: fixed (Fig. 3E, F) or unfixed (Fig. 3G, H). Partitioning the data sets revealed potential variation in the RTA-DNA interactions, whereas fixation preserves all interactions at a given time, only RTA-DNA interactions capable of surviving column purification would be maintained in unfixed conditions. Analysis of RTA-OriLyt-L binding location revealed a high frequency of RTA mapped to 900-1000 bp region, however several additional peaks occurred at 1200-1300, 1500- 1600 bp in the unfixed conditions (Fig. 3D). These sites coincide with AP-1 sites (green line) and AT-rich regions (cyan line). The differential binding peaks are summarized in Table 2.

**Figure 3.**
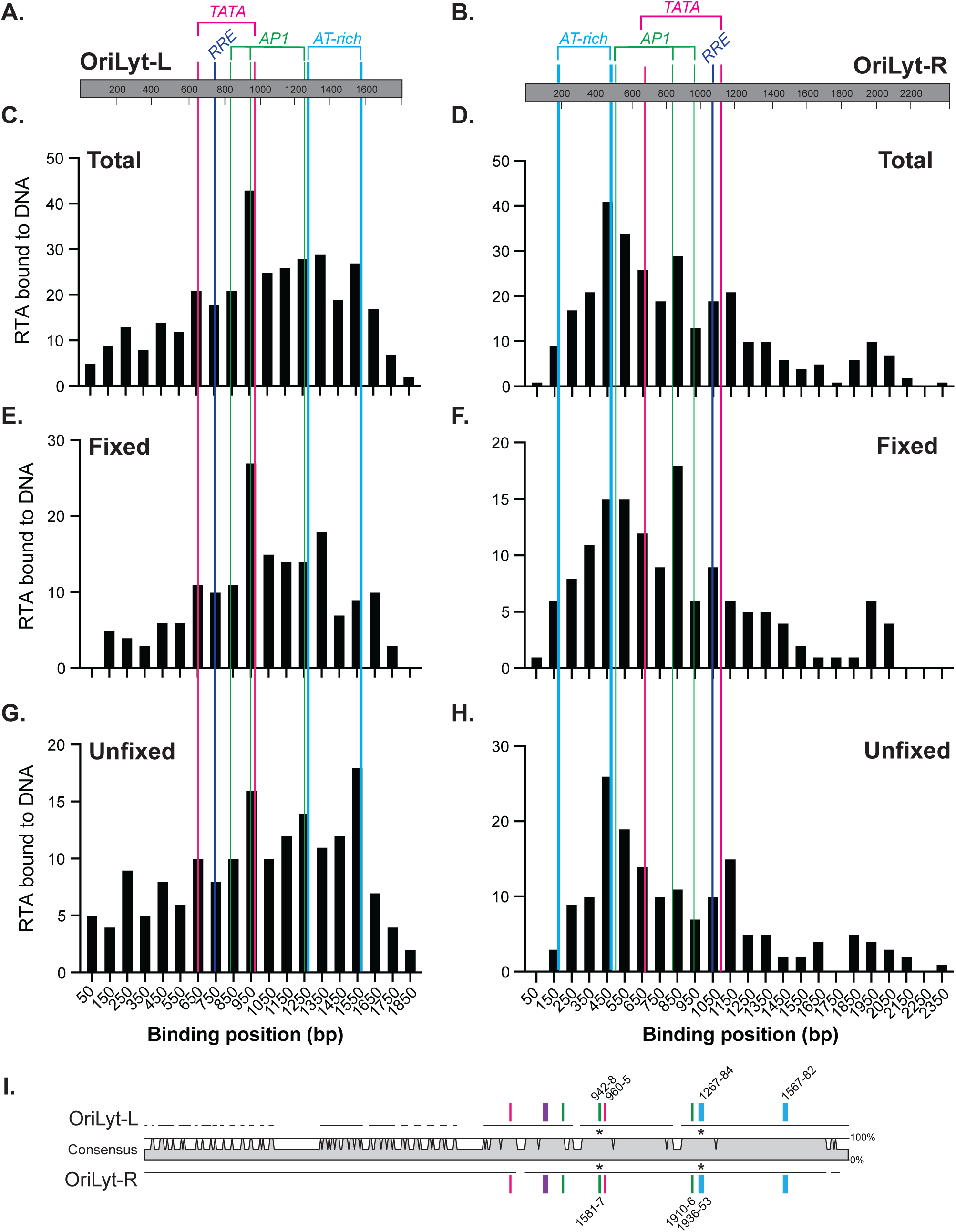
Corresponding base pair binding position of RTA bound to DNA lytic origin sequences. A-B. Schematic of OriLyt-L and OriLyt-R including relative position of key DNA sequences AP-1 (green), AT-rich (cyan), RRE (purple), TATA-box (magenta). C-H. Histograms of the number of RTA molecules bound to KSHV OriLyt. X-axis shows binding position (base pair, bp) bin center value, +/25bps. Y-axis frequency of protein count. Total protein frequencies for C. OriLyt-L, or D. OriLyt-R compared to the E-F. fixed and G-H. unfixed protein-DNA sample preparations. I. Sequence alignment homologous regions of OriLyt-L and -R. Black lines indicate origin DNA sequences and shows gaps in alignments. The consensus from 0 to 100% as compared between two origin sequences, grey shading to 100% indicates identical nucleotide sequence at given location. Nucleotide position indicated for regions with highest frequency peaks for RTA bound to each OriLyt and asterisks (*) indicates identical region in both OriLyt-L and -R.

**Table 2.** High frequency RTA binding locations (bp)

| **Table 2. High frequency RTA binding locations (bp)** |  |  |  |
| --- | --- | --- | --- |
| Origin sequence (SA-end to RTA) | Primary 1 peak | Peak 2 | Peak 3 |
| **OriLyt-L** |  |  |  |
| Total (n= 334 RTA proteins) | 900-1000 |  |  |
| Fixed (n= 171 RTA proteins) | 900-1000 |  |  |
| Unfixed (n= 163 RTA proteins) | 1500-1600 | 900-1000 | 1200-1300 |
| **OriLyt-R** |  |  |  |
| Total (n= 292 RTA proteins) | 400-500 |  |  |
| Fixed (n= 138 RTA proteins) | 800-900 | 400-600 |  |
| Unfixed (n= 154 RTA proteins) | 400-500 |  |  |

The most abundant binding site for RTA-OriLyt-R was the region between 400-500 bp from the SA-labeled end of the DNA. This region corresponds with an AP-1 site and an AT-rich region (Fig. 3D, cyan and green lines). We observed additional high occupancy RTA binding sites when the data was separated into fixed and unfixed groups. For fixed samples, the greatest number or RTA molecules bound to the region between 800-900 bp; coinciding with an AP-1 site (Fig. 3G, green line), however the 400-500 bp region was also a high abundance protein binding location, similar observation to unfixed (Fig. 3H). In the unfixed samples, a third, less prominent peak emerged at 1100-1200 bp; this region is in close proximity to the RTA RRE (Fig. 3H, magenta line).

An additional way to compare RTA binding preference is to examine conserved similarities and differences between the two OriLyt sequences. As previously described, OriLyt-L and -R contain homologous sequences (44). An alignment of the reverse complementary sequence of OriLyt-L aligned with OriLyt-R (Fig. 3I, 100% homology full grey shading) shows the sequence similarity and distribution of known protein binding consensus sequences as indicated by colored lines. The relative base pair locations for each sequence are indicated, as well as the common binding sites measured in Figure 3C-H, (Fig. 3I, asterisks). The relative bp for the complement and reverse complement sequences are listed in Supplemental Table S3. The highest peaks for both OriLyt-L and -R overlapped at two locations and include AP-1 (green bar) and AT-rich regions (blue bar). When we compare OriLyt-L and -R there are several regions, Fragments (Fr) A-C with different RTA occupancy, these observations were further analyzed using gel-based methods.

### Complementary EMSA further defined protein binding sequences

EMSA complemented our TEM studies and confirmed RTA binding to several AP-1 sequences (Fig. 4). The protein binding regions identified using TEM as high frequency were amplified or synthesized. Due to the highly repetitious stretches of A-T and G-C, PCR primer design proved challenging and an alternative approach using annealed synthesized single stranded DNA oligos was needed to generate the smaller DNA fragments (99-239 bp). These DNA lengths are at the lower level of TEM resolution. Figure 4A maps the sections of highest homology and smaller OriLyt fragments (FrA, B, C tan boxes). OriLyt-L and -R FrA includes the known RTA RRE (ATGGGTGGCTAAC,(39) purple) and AP1-1 (green). OriLyt-R FrA includes known RTA RRE (purple) and AP1-1 (green) and a nearly identical second RTA RRE (A<u>C</u>G<u>CT</u>TGGCTAAC, purple) that is 3 nucleotides different from known RRE. Fragment B, FrB, contains the conserved AT-rich and AP1-3. OriLyt-L fragment C, FrC, includes AP1-2 and the AT-rich region. The relative bp for the complement and reverse complement fragment sequences are listed in Supplemental Table S3.

**Figure 4.**
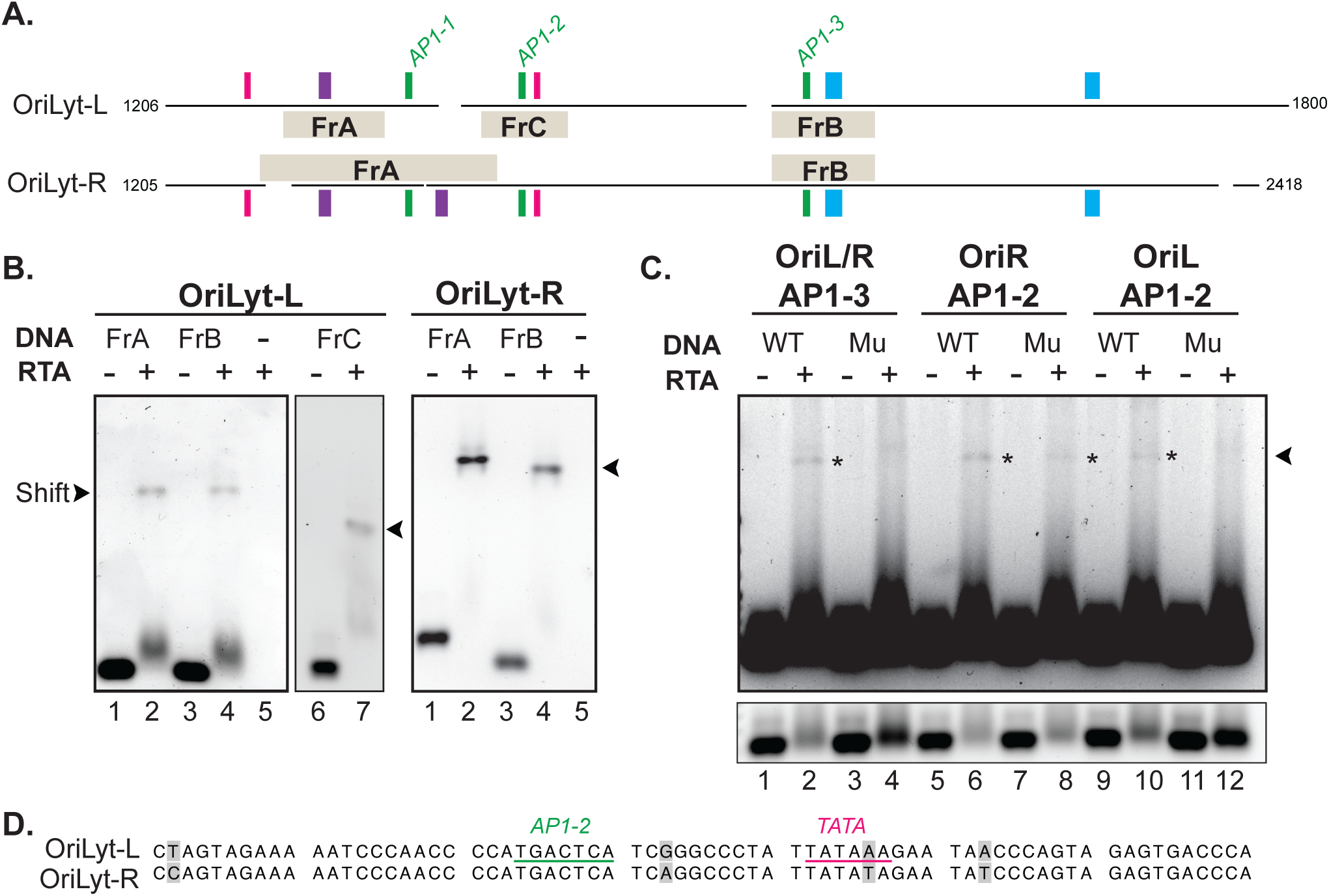
EMSA analysis of OriLyt-L and -R fragments and AP-1 sites. A. Schematic of OriLyt-L and OriLyt-R including relative position of key DNA sequences: fragments A-C (tan), AP-1 (green), AT-rich (cyan), RRE (purple), TATA-box (magenta). B. Representative EMSA agarose gel of OriLyt-L, fragment A (FrA) and fragment B (FrB) and -R FrA and FrB in the absence (lanes 1, 3) or presence (lanes 2, 4) of RTA. C. Representative EMSA agarose gel of OriLyt mutagenesis analysis of AP-1 sites. OriLyt-L and -R wild-type (WT) and mutant (Mu) AP1-3 and AP1-2 in the absence (lanes 1, 3, 5, 7, 9, and 11) or presence (lanes 2, 4, 6, 8, 10 and 12) of RTA. Shifted DNA indicated by asterisks (*lanes 2, 6, 8, 10). D. Sequence alignment of OriLyt-L and -R wild-type region surrounding AP1-2 (green line) and TATA-box (pink line), grey boxes indicate different nucleotides.

The OriLyt-fragments shifted to a higher molecular weight band (arrowhead) when incubated with RTA (Fig. 4B, OriLyt-L lanes 2, 4 and 7 and OriLyt-R lanes 2 and 4). To confirm and refine the RTA binding sequence short DNA oligos (99 nucleotides) containing OriLyt-AP1 sites (1–3) and mutant AP-1 sites, TG<u>C</u>CT<u>T</u>A,(52) were synthesized and annealed for EMSA (Supplemental Table S2). The AP-1 site containing oligos incubated with RTA shifted (Fig. 4C Lanes 2, 6, and 10). Mutant AP1-3 (lane 4) and OriLyt-L AP1-2 (lane 12) oligos incubated in the presence of RTA did not produce a shifted band. The absence of the shifted band was confirmed by the intensity of the free DNA (Fig. 4C, lower panel, shorter exposure; lanes 2 and 10 compared to 4 and 12, respectively). The mutation of AP1-2 in OriLyt-R did not diminish the shifted band indicative of RTA binding (Fig. 4C, lane 8). This data and the comparatively small, high protein frequency peak in unfixed (Fig. 3H) OriLyt-R at the corresponding region suggests RTA does not specifically bind to OriLyt-R AP1-2. However, the AP1-2 region within OriLyt-L maintained a high frequency of RTA binding in both fixed and unfixed samples Fig. 3E, G).

Additional sequence analysis identified five nucleotide differences between OriLyt-L and -R in the area surrounding AP1-2 (Fig. 4D, shaded boxes).

### Comparison of the protein area of RTA with protein standards

While we used positional data to approximate regions where RTA bound to viral DNA, we also measured the 2D projected area of each protein in the micrographs to approximate the oligomeric state of RTA under different conditions (Fig. 5). We observed RTA with a range of sizes when bound to OriLyt. RTA was compared with protein standards conalbumin and alcohol dehydrogenase to approximate the oligomeric state of RTA (Fig. 5A-B). RTA has been previously shown as a dimer or higher order multimer when binding to viral promoters and studies have suggested that RTA binds to the OriLyt as a dimer (19,53). The measured area (nm²) of RTA, streptavidin, and protein standards (conalbumin and alcohol dehydrogenase) were compared (Fig. 5B). Analysis of variance between the populations of molecular standards confirmed that differences of the means between streptavidin, conalbumin and alcohol dehydrogenase were statistically distinct groups. Likewise, the difference between the area of full-length RTA bound to Orilyt-R and unbound truncated ORF50/RTA was found to be statistically significant, demonstrating the sensitivity of our assay (Fig. 5B).

**Figure 5.**
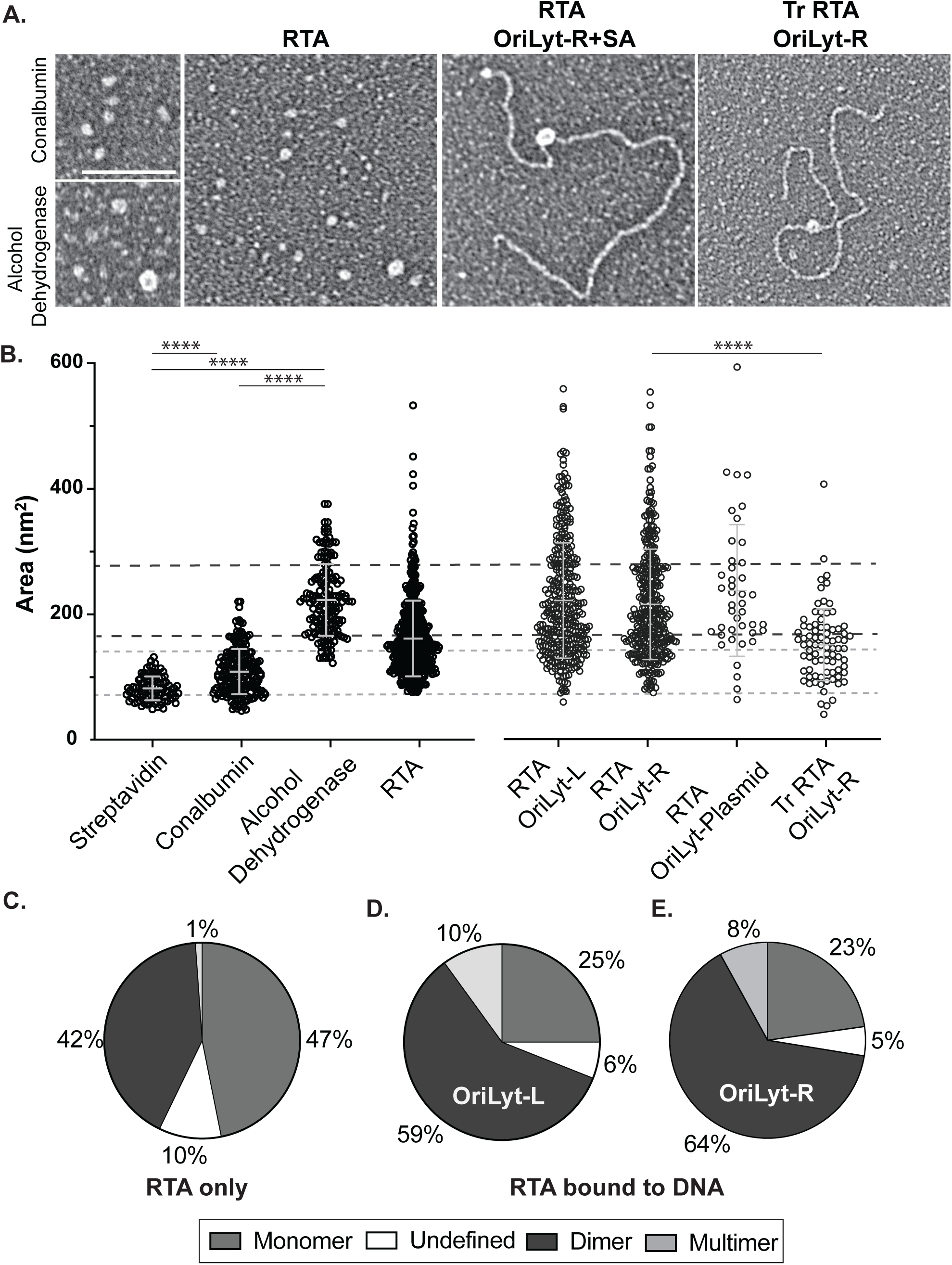
Protein size comparison of RTA in absence and presence of OriLyt DNA. A. Representative micrographs of conalbumin, alcohol dehydrogenase, RTA with or without OriLyt and truncated RTA with OriLyt. Scale bar = 200 nm. B. Dot plot of the area (nm^2^) of each protein: streptavidin (n=101), conalbumin (n=200), alcohol dehydrogenase (n=163), RTA (n=458), RTA bound to OriLyt-L (n=334), RTA bound to OriLyt-R (n=292), RTA bound to OriLyt-plasmid (n=42) and truncated RTA bound to OriLyt-R (n=81) with the mean+/-SD. Dashed line indicates standard deviation of conalbumin (grey line) or alcohol dehydrogenase (black line). Statistical significance by unpaired t-test, p<0.0001 denoted by ****). C-E. Pie charts showing the percent of RTA monomers, dimer or multimers in the absence or presence of Orilyt-L or -R.

Conalbumin and alcohol dehydrogenase have molecular weights that coincide with the predicted molecular weights of the monomeric (75 kDa) and dimeric (150 kDa) forms of RTA. The average area for measured conalbumin was 109± 36 nm², which established a baseline for RTA monomers (Fig. 5B). The average area for measured alcohol dehydrogenase was 223 ±57nm² was used as the approximate RTA dimer (Fig. 5B). RTA proteins measured area between 145-166 nm² were excluded as either monomeric or dimeric in our investigations and were denoted as undefined. Based on these parameters, the RTA protein alone was observed as a monomer and dimer at almost equal proportion, (42% and 47%, Fig. 5C). In contrast, when RTA was bound to OriLyt-L, 25% of measured protein were in the range for monomeric conformation, 59% dimeric and 10% were multimeric (Fig. 5D). Similarly, when RTA was bound to the OriLyt-R 25% of were monomeric, 64% dimeric and 23% were multimeric (Fig. 5E). Indicating the RTA preferential binds to lytic origin DNA as a dimer.

### Characterizing the DNA binding site of protein monomers and dimers

To determine where our defined RTA monomers and dimers bind OriLyt, we examined the relationship between binding position frequencies and protein conformation concurrently (Fig. 6). Schematics of OriLyt-L and -R denote the DNA domains (RRE, AP-1, TATA, AT-rich, Fig. 6A-B). When comparing RTA binding tendencies in terms of protein monomers and dimers we observed conserved patterns of dimeric RTA and differential binding behavior of monomeric RTA, when comparing OriLyt-L and -R. For both origin sequences, RTA dimers localized at highest frequency at the AP1-3/AT-rich regions (Fig. 5C, 1201-1400bp and 5D, 401-600bp). However, analysis of monomeric RTA proteins shows a high proportion of RTA monomers bound to two OriLyt-R regions, one coinciding with the dimer peak and the other with previously predicted RRE and TATA box region (Fig. 5F, 1001-1200bp), while RTA monomers bound to OriLyt-L bind throughout the DNA sequence (Fig. 5E).

**Figure 6.**
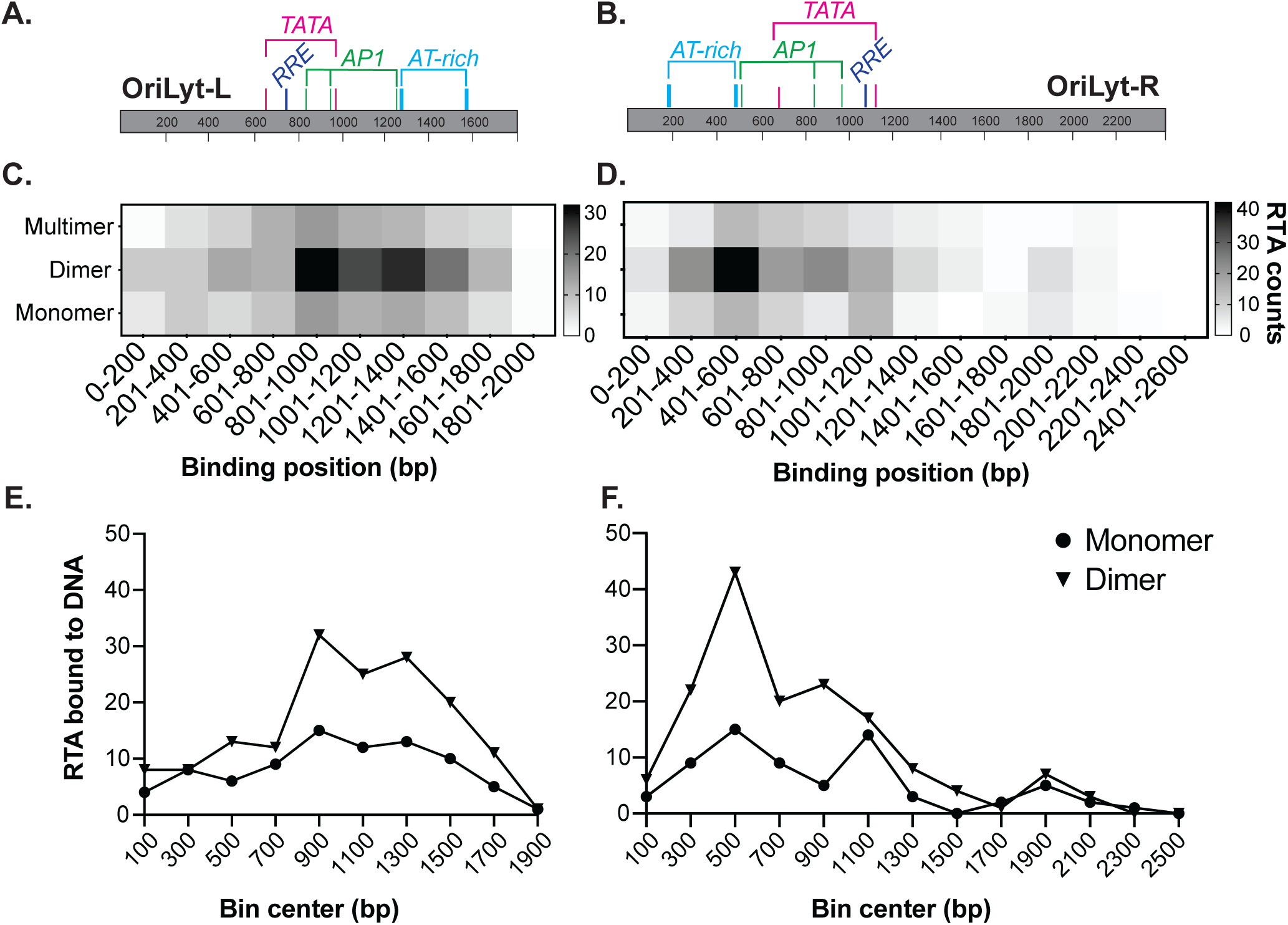
Combinational analysis of protein size and DNA binding location. A-B. Schematic of OriLyt-L and -R including relative position of key DNA sequences AP-1 (green), AT-rich (cyan), RRE (purple), TATA-box (magenta). C-D. Heat maps of RTA monomers, dimers or multimers bound to OriLyt-L and -R. E-F. Graphs of the bin center of RTA monomers and dimers bound to OriLyt-L and -R.

Together these analyses of TEM micrographs revealed RTA dimers have a propensity to bind to more specific DNA binding locations while monomers may have less of a binding sequence specificity. The data can be used to possible infer the mechanism of RTA binding. Potentially, RTA monomers scan the DNA until a second RTA protein binds and increases binding specificity to a specific DNA site. Furthermore, in the majority of looped DNA associated with RTA (approximately 30 molecules) we measured RTA as a dimer or multimer (data not shown). This observation indicates more than one RTA proteins binding to separate DNA sites while simultaneously maintaining protein-protein interactions.

## Discussion

In this study we have provided data using a microscopy based-method for examining purified protein behavior at the molecular level, expanded upon our understanding of RTA DNA-binding position and identify the oligomeric structure of RTA during origin DNA binding. We demonstrated that RTA bound to discreet regions of OriLyt DNA with high frequency (Fig. 3-4) and does so primarily as a dimer (Fig. 5). Our data supports what has been previously posited about RTA oligomeric state using traditional methods such as EMSA, SDS-PAGE and immunoblotting (36,53). However, our observations of RTA’s heterogenous oligomeric state when bound to viral DNA compared to protein alone expanded upon the understanding by showing direct visual evidence via a primarily TEM approach.

By comparing RTA alone and RTA bound to DNA, we determined that the formation of larger size multimers is associated with protein-DNA binding. Interestingly, our data supports that the regions where RTA binds with greatest abundance do not correlate with known RTA response elements (54). We observed the most frequent binding were correlated to the position of AP-1 sites, TATA boxes and/or A-T rich regions (Fig. 3I). This pattern was consistent between the OriLyt-L and -R. OriLyt and RTA aggregation at these sites, may play a role in recruitment of the core viral DNA replication machinery(41,55) to AT-rich regions (36), or other transcription factors to AP-1 and TATA box locations during the KSHV lytic phase (33).

Further, these data support the idea that RTA may play dual, if not several roles in the KSHV lytic cycle, and the differential binding we observed may be altered by the presence of additional viral or cellular proteins. A limitation of our purified system is the inability to effectively replicate the cellular environment of viral DNA replication in the nucleus. Several previous studies characterizing RTA interactions with DNA used whole cell lysates and generated conclusions about RTA from cellular extracts, which reports either average or focuses on the species at greatest abundance in a population. This study’s approach using TEM and downstream quantification is unique in that we directly visualized and measured hundreds of individual RTA proteins and RTA bound to OriLyt DNA.

Beyond facilitating the analysis of heterogeneous populations, TEM is advantageous for reasons including: (i) TEM uses highly purified components interacting in solution before being preserved for TEM visualization, allowing for real-time capture of protein-DNA interactions, (ii) TEM is compatible with an array of DNA lengths, that do not need to be affixed to the surface as is the case for other high-resolution microscopic approaches, such as atomic force microscopy (ATM). Our future studies will seek to improve our system by standardizing sample preparation and streamlining the data analysis process to more efficiently characterize protein-DNA interactions. Additionally, immunogold labeling(52) and the use of new technologies will allow for differentiation of individual species on the TEM grid by photosensitizing fluorescent tracers or oxidation, can play a critical role in further analyzing multiple proteins simultaneously using TEM (56). Likewise, the potential to ‘color-code’ RTA isoforms on the TEM grid could lead to new discoveries about the sub-structure of the protein and how that impacts protein behavior. Our long-term goal is to examine proteins in concert to determine how protein-protein interactions may alter DNA-protein binding specificity.

## Acknowledgements

This work was supported by the following NIH grants: K12-GM000678, SC2GM136527 and P01CA019014. We thank Dirk Dittmer, Ryan McNamara, Smaranda Wilcox for expert advice and past and present members of the Costantini laboratory for scientific discussions. We dedicate this work to Roger Ibis for generously donating time and technical expertise to relocate the transmission electron microscope.

## Supplemental Figure legends

**Supplemental Figure S1**. Optimization of TEM DNA-protein data acquisition and analysis. A. Schematic of binding reaction for purified proteins and target DNA OriLyt. B. Representative TEM micrographs of RTA bound (white arrowhead) to super twisted and relaxed plasmid DNA containing OriLyt. C. Schematic of binding reaction for purified proteins with SA end-labeled-DNA and column fractionation. D. Representative TEM micrograph of RTA bound (white arrowhead) to streptavidin (black arrowhead), SA-end labeled OriLyt fragment DNA in various conformations. Scale bar=200nm.

**Supplemental Figure S2**. OriLyt biotin incorporation and streptavidin end-labeling. A. Schematic of OriLyt-L containing plasmid with restriction enzyme sites (HindIII/EcoRI) and location of dCTP-biotin incorporation. B. Representative micrographs of OriLyt-L in absence or presence of streptavidin (black arrowhead). A. Schematic of OriLyt-R containing plasmid with restriction enzyme sites (NdeI/XhoI/ScaI) and location of dCTP-biotin incorporation. B. Representative micrographs of OriLyt-R in absence or presence of streptavidin (black arrowhead). Scale bar=200nm.

**Supplemental Table S1.**
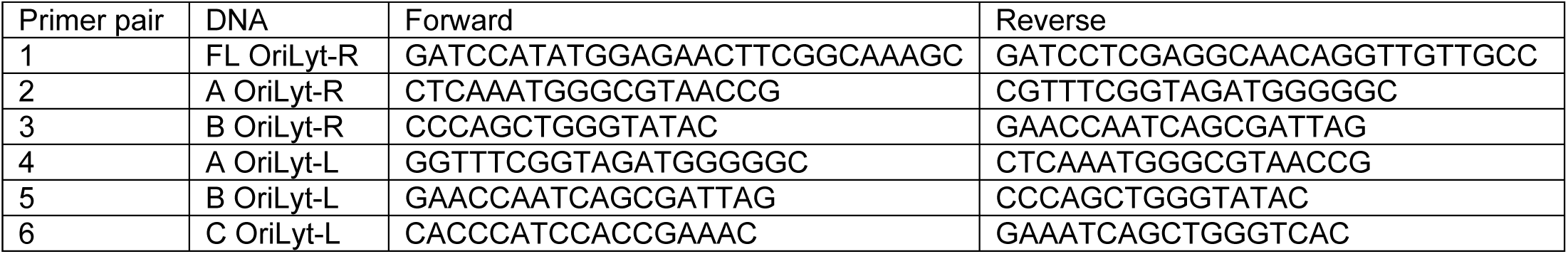
PCR primers.

**Supplemental Table S2.**
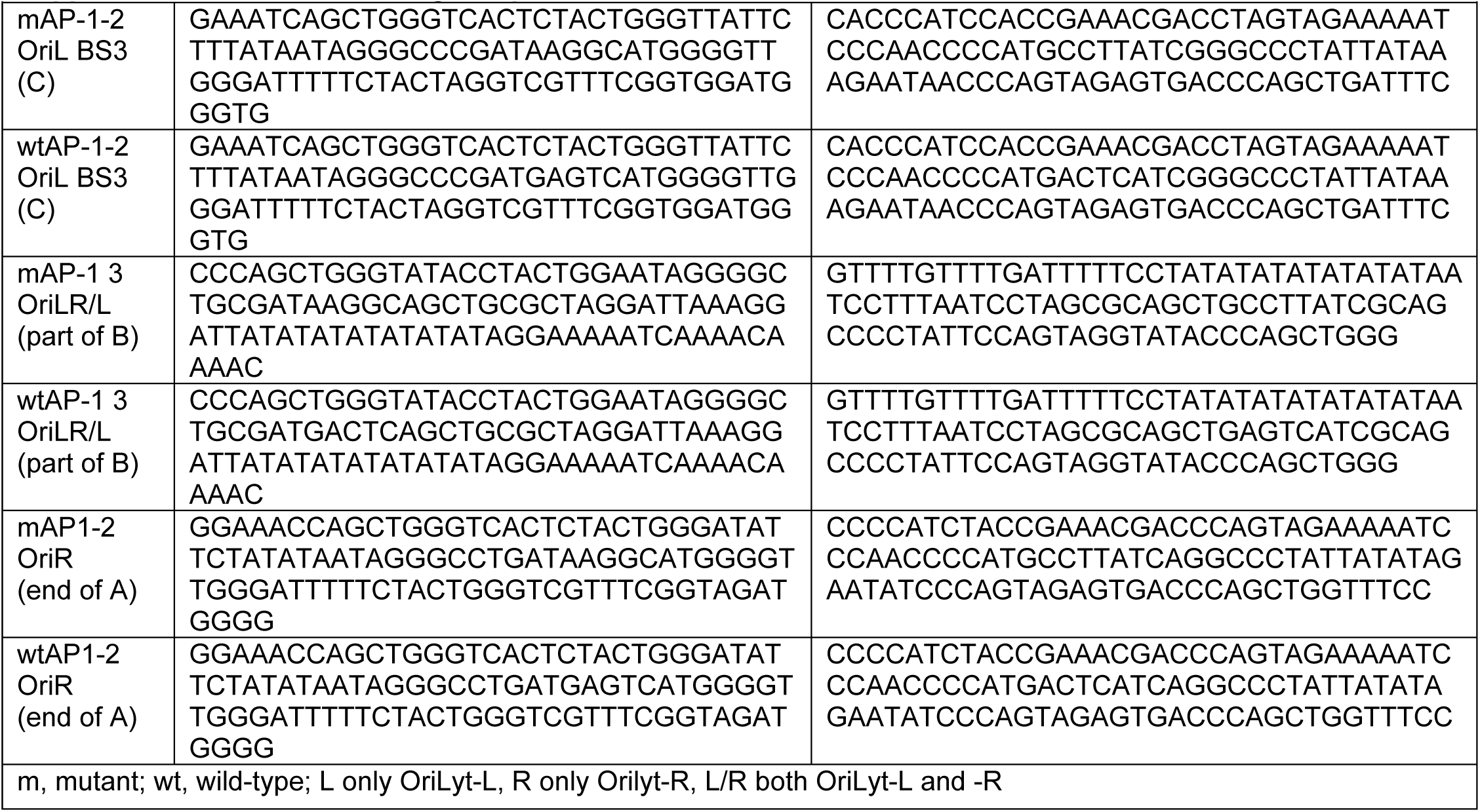
DNA oligo sequences.

**Supplemental Table S3.** Summary of KSHV lytic origin DNA sequences.

| **OriLyt-R** | **Unlabeled** | **SA-labeled** |
| --- | --- | --- |
| RTA RRE | 1354-1366 (13bp) | 1053-1065 |
| FrA | 1316-1554 (239bp) | 865-1103 |
| RTA RRE 2 | 1485-1497 (13bp) | 922-934 |
| FrB | 1874-1991 (118bp) | 428-545 |
| AP1-3 | 1910-1916 (7bp) | 503-509 |
| AP1-2 | 1581-1587 (7bp) | 832-838 |
| AP1-1 | 1453-1459 (7bp) | 960-966 |
| TATA | 1765-1770 (6bp) | 649-654 |
| TATA | 1298-1303 (6bp) | 1116-1121 |
| AT rich | 2236-2251 (16bp) | 168-183 |
| AT rich | 1936-1953 (18bp) | 466-483 |
| **OriLyt-L** | **Unlabeled** | **SA-labeled** |
| RTA RRE | 1051-1063 (13bp) | 738-750 |
| FrA | 990-1104 (115bp) | 697-811 |
| FrB | 479-596 (118bp) | 1205-1322 |
| FrC | 804-902 (99bp) | 899-997 |
| AP1-3 | 554-560 (7bp) | 1241-1247 |
| AP1-2 | 853-859 (7bp) | 942-948 |
| AP1-1 | 958-964 (7bp) | 837-843 |
| TATA | 836-841 (6bp) | 960-965 |
| TATA | 1144-1149 (6bp) | 652-657 |
| AT rich | 219-234 (16bp) | 1567-1582 |
| AT rich | 517-534 (18bp) | 1267-1284 |

